# Reduced sleep pressure in young children with autism

**DOI:** 10.1101/706135

**Authors:** Ayelet Arazi, Gal Meiri, Dor Danan, Analya Michaelovski, Hagit Flusser, Idan Menashe, Ariel Tarasiuk, Ilan Dinstein

## Abstract

**Study Objectives:** Sleep disturbances and insomnia are highly prevalent in children with Autism Spectrum Disorder (ASD). Sleep homeostasis, a fundamental mechanism of sleep regulation that generates pressure to sleep as a function of wakefulness, has not been studied in children with ASD so far, and its potential contribution to their sleep disturbances remains unknown. Here, we examined whether slow wave activity (SWA), a measure that is indicative of sleep pressure, differs in children with ASD.

**Methods:** In this case-control study, we compared overnight electroencephalogram (EEG) recordings that were performed during Polysomnography (PSG) evaluations of 29 children with ASD and 23 typically developing children.

**Results:** Children with ASD exhibited significantly weaker SWA power, shallower SWA slopes, and a decreased proportion of slow wave sleep in comparison to controls. This difference was largest during the first two hours following sleep onset and decreased gradually thereafter. Furthermore, SWA power of children with ASD was significantly, negatively correlated with the time of their sleep onset in the lab and at home, as reported by parents.

**Conclusions:** These results suggest that children with ASD may have a dysregulation of sleep homeostasis that is manifested in reduced sleep pressure. The extent of this dysregulation in individual children was apparent in the amplitude of their SWA power, which was indicative of the severity of their individual sleep disturbances. We, therefore, suggest that disrupted homeostatic sleep regulation may contribute to sleep disturbances in children with ASD.

**Statement of significance:** Sleep disturbances are apparent in 40-80% of children with autism. Homeostatic sleep regulation, a mechanism that increases the pressure to sleep as a function of prior wakefulness, has not been studied in children with autism. Here, we compared Polysomnography exams of 29 children with autism and 23 matched controls. We found that children with autism exhibited reduced slow-wave-activity power and shallower slopes, particularly during the first two hours of sleep. This suggests that they develop less pressure to sleep. Furthermore, the reduction in slow-wave-activity was associated with the severity of sleep disturbances as observed in the laboratory and as reported by parents. We, therefore, suggest that disrupted homeostatic sleep regulation may contribute to sleep disturbances of children with autism.

## Introduction

Autism Spectrum Disorders (ASD) are a family of heterogeneous neurodevelopmental disorders characterized by impairments in social interaction and by restricted interests and repetitive behaviors ^1^. Sleep disturbances appear in 40-80% of children with ASD^2–5^, as compared with 20-40% of typically developing children^6,7^. Symptoms include prolonged sleep latency, shorter sleep duration, and increased wake periods during the night, as reported by both subjective parental questionnaires and actigraphy measures^8–14^. Poor sleep in children with ASD is associated with increased sensory sensitivities^15–18^ and increased aberrant behaviors^18–21^, which impair the quality of life of affected families^14^. PSG studies have corroborated the existence of sleep disturbances in children with ASD^22–28^, but have yielded mixed results regarding potential abnormalities in sleep architecture. While some have reported that children with ASD exhibit decreased slow wave sleep (SWS, stage N3)^22–25^ or rapid eye movement (REM)^11,22,28^ durations, others have not^23,26–28^.

It has been proposed that the sleep disturbances of children with ASD are caused by an interaction of several behavioral and physiological factors including anxiety^21^, poor sleep hygiene^29^, sensory hyper-sensitivities^15^, abnormalities with the melatonin system (i.e., circadian rhythm)^30^, and obstructive sleep apnea (OSA)^31^. To date, the potential contribution of disrupted sleep homeostasis to the emergence of insomnia in children with ASD has not been examined.

Sleep homeostasis is a critical mechanism of sleep regulation that increases the pressure to sleep as a function of time spent awake^32,33^. Larger sleep pressure generates deeper slow wave sleep, which can be quantified by the power of slow wave activity (SWA, EEG power in the Delta band, 0.75-4 Hz)^34^. Deep sleep is essential for proper cognitive function^35^, stabilizing synaptic plasticity^36,37^, and enabling learning and memory consolidation^38^. To our knowledge, only three studies to date have quantified the amplitude of SWA in ASD and all were performed with high-functioning adolescents and adults^39–41^. Two of these studies reported significantly weaker SWA in the ASD group^39,40^, a difference that was particularly large during the first 2-3 hours of sleep.

To evaluate potential impairments in the SWA of children with ASD, we examined PSG recordings from 29 children with ASD and 23 typically developing controls. We quantified SWA power, SWA slope, and traditional sleep staging in 1-hour segments from sleep onset, and assessed whether significant differences were apparent across groups during specific segments of sleep. In addition, we examined whether individual differences in SWA could explain differences in the severity of sleep disturbances as observed in the sleep laboratory and as reported by the parents at home.

## Methods

### Subjects

We recruited 34 children with ASD (9 females), mean age 4.6 years old (range 1.9-7.8), from the National Autism Research Center of Israel^42^, which is part of Ben Gurion University of the Negev, and located in Soroka University Medical Center (SUMC). Approximately 200 children are referred to SUMC from the community (i.e., education system, primary care, etc.) annually with a suspicion of ASD and approximately 80% of these children receive a positive diagnosis. We randomly approached 70 families who received a positive diagnosis between April 2017 to August 2018, and 34 of them agreed to participate in the study. Families were recruited regardless of specific symptom severities. Informed consent was obtained from all parents of ASD children, who were reimbursed for their participation.

All children with ASD were referred by the research team to a PSG evaluation at the Sleep-Wake Disorder Unit of SUMC. We excluded five children with ASD from the final analysis due to poor PSG quality (n=4) or evidence of Obstructive Sleep Apnea (OSA, n=1).

All children with ASD were diagnosed with autism, independently, by a physician and a developmental psychologist according to DSM-5 ^1^ criteria. Of the 29 children in the final analysis, 26 children with ASD also completed the autism diagnostic observation schedule (ADOS)^43^, the remaining 3 children did not complete the ADOS due to lack of availability. Mean ADOS scores were: Social Affect 14.9±5, Restricted and Repetitive Behaviors 4.3±1.7, and total score 19.2±6.1. Parents of 27 children with ASD completed the Hebrew version of the child sleep habit questionnaire (CSHQ)^44,45^. Parents of 2 additional children did not complete the questionnaire. CSHQ scores were compared to the mean scores of a large cohort of typically developing children without sleep problems^44^. The biggest sleep concerns for the ASD children involved bedtime resistance, sleep anxiety, and excessive daytime sleepiness (Table 1).

**Table 1:**
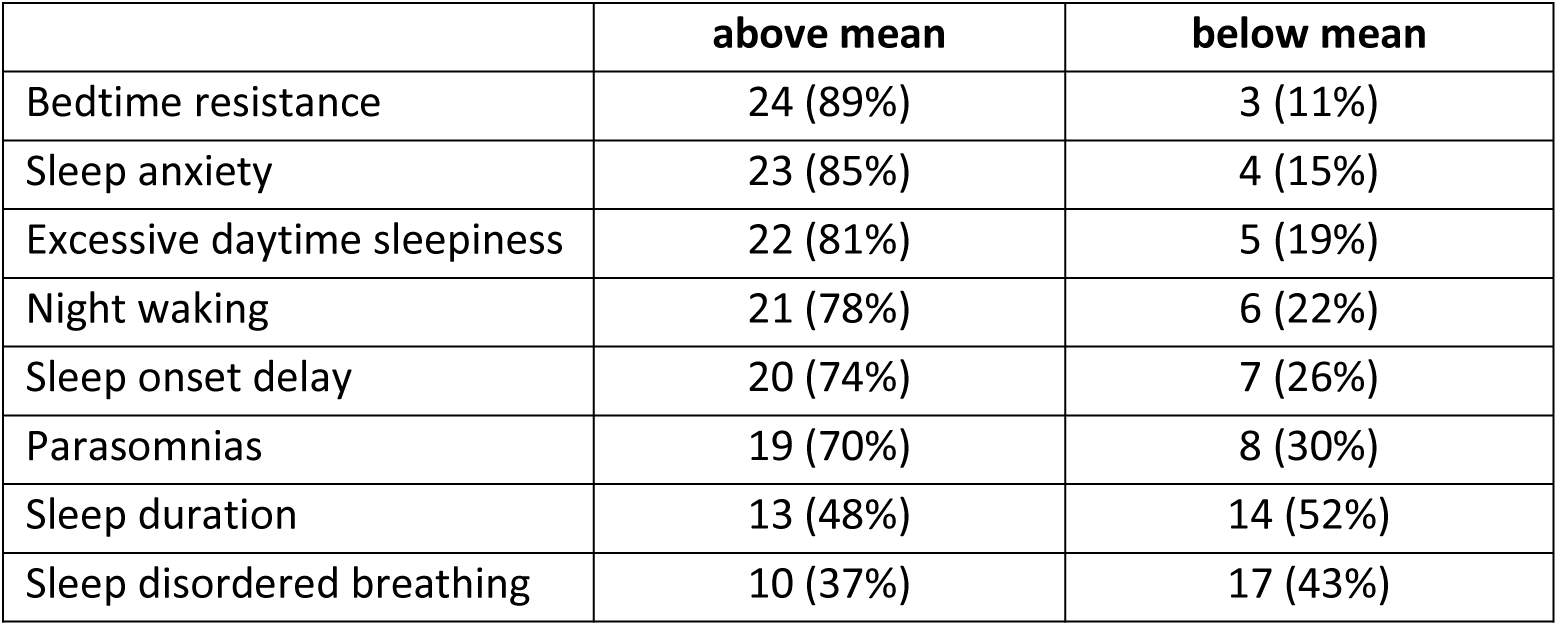
sleep concerns in children with autism. Sleep concerns as reported by parents of the 27 children with ASD who completed the CSHQ. The CSHQ scores sleep disturbances in 8 domains. We present the number of ASD cases (and percentage) who had scores that were above/below the mean score of previously published CSHQ scores from a large population of typically developing children^44^.

The control group included 23 children who were identified retrospectively from children who were referred to PSG evaluation at the same SUMC unit and their discharge letter indicated that they did not have any clinical findings. Reasons for initial referral of these children included snoring (n = 18) or other sleep problems such as bedwetting and daytime sleepiness (n = 5). All of the control children were screened negative for neurological, psychiatric, or developmental disorders, using a detailed clinical history questionnaire that was completed by the parents.

The Helsinki committee at SUMC approved this study, which was carried out under the guidelines of the Helsinki declaration.

### Polysomnography

All parents were instructed to keep a regular sleep-wake schedule on the day of the PSG evaluation. The PSG study started at 8:30PM and ended at 6:00AM on the following morning. Children were connected to a clinical PSG system (SomniPro 19 PSG, Deymed Diagnostic, Hronov, Czech Republic) by a technician with over 5 years of experience. All participants were connected to six EEG electrodes, (C3, C4, O1, O2, A1, and A2 according to the international 10-20 system; sampling frequency: 128Hz; resolution: 16 bit), EOG, EMG and ECG electrodes, abdomen and chest effort belts to measure respiratory activity, and an oxygen saturation sensor. In some cases, where the child did not cooperate, this procedure was completed after the child fell asleep, typically within 10 minutes of sleep onset. Four children with ASD were being treated with Melatonin, but did not take Melatonin on the day of the exam. Derivations C3/A2 and C4/A1 were used for sleep-stage scoring, which was determined blindly by one of the investigators (AT) according to the American Academy of Sleep Medicine criteria^46^. An Apnea Hypopnea Index (AHI) was calculated as the number of respiratory events resulting in either arousal or oxygen desaturation of >4%, per hour of sleep^47^.

### EEG analysis

#### Preprocessing

Data were analyzed offline using MATLAB (*Mathworks Inc. USA*) and the EEGLAB toolbox^48^. EEG data was re-referenced to the bilateral mastoids, filtered using a 0.75Hz FIR high-pass filter (cutoff frequency at −6db: 0.37Hz, transition band width: 0.75Hz) and 20Hz FIR low-pass filter (cutoff frequency at-6db: 22.5, transition band width: 5Hz), and then divided into consecutive 30-sec epochs. Epochs with manually identified artifacts such as movement or muscles contractions were removed (mean percentage of removed epochs in the ASD group: 18.9%, and in controls: 16.5%).

#### EEG data analysis

Each 30 second epoch was subdivided into consecutive 4 second segments with a 2 second overlap, using a hamming window. The power spectrum was computed for each 4 second segment using the FFT function as implemented in MATLAB (frequency resolution of 0.25 Hz), and then averaged across segments of each 30 second epoch. Absolute power was calculated for the Delta (1-4Hz), Theta (4-8Hz), Alpha (8-13Hz), and Beta (13-20Hz) frequency bands as the sum of power across these frequencies.

To compute the slope of SWA we re-filtered the EEG data using a 0.75-4Hz band-pass filter. Slow waves were identified in each 30 second epoch as negative peaks, with subsequent zero-crossing, that were separated by 0.25-1 sec. We calculated the slope of each wave as the amplitude of the negative peak divided by time to the next zero crossing (i.e., ascending slope) and then computed the mean slope across all waves in each 30 second epoch. Descending slopes were also computed, as the negative peak divided by the time to the previous zero crossing. Both ascending and descending slope measures yielded equivalent results and only ascending slopes are reported in the manuscript.

### Statistical analysis

Statistical analysis was performed using MATLAB (*Mathworks Inc. USA*). SWA power, SWA slope, or the percentage of sleep at each sleep stage (i.e., N2, N3 or REM) were analyzed using 2-way ANOVAs with group as one factor and time (i.e. hour since sleep onset) as a second factor. Additional comparisons across ASD and control groups were performed using two-tailed t-tests with unequal variance. We also computed the effect size for each comparison using Cohen’s d. When performing comparisons across groups for each frequency or each 1-hour segment of the night (from sleep onset), we used the false discovery rate (FDR) correction^49^ to control for the multiple comparisons problem. The relationship between CSHQ scores and EEG power was assessed using Pearson’s correlations. The statistical significance of the correlation coefficients was tested with a randomization test where we shuffled the labels of the subjects before computing the correlation. We performed this procedure 10,000 times to generate a null distribution for each relationship and assessed whether the true correlation coefficient was higher than the 97.5^th^ percentile or lower than the 2.5^th^ percentile of this null distribution (equivalent to p=0.05 in a two-tailed t-test).

## Results

Children with ASD spent significantly less time in bed, and their total sleep time was shorter than that of controls (Table 2). Sleep efficiency, percentage of wake after sleep onset (WASO), sleep latency, and arousal index did not differ across groups. The mean apnea-hypopnea index (AHI) was <1 in both groups, demonstrating that none of the participating children had OSA. Note that ~50% of the ASD children and 17% of the controls were connected to the PSG apparatus only after sleep onset. As a result, these PSG recordings began approximately 10 minutes after the children fell asleep. This is a common issue in PSG studies with ASD children, where measures of time in bed, sleep latency, and sleep efficiency are often distorted due to poor cooperation at the initiation of the exam.

Small differences in sleep architecture across groups were found when examining the proportion of sleep stages throughout the entire night. Children with ASD exhibited a significantly higher percentage of N2 sleep (p=0.03, Cohen’s d=0.61; Table 2), and a marginally significant reduction in the percentage of REM sleep (p=0.06, Cohen’s d=0.54). Larger differences across groups were revealed when dividing the sleep period of each child into two equal halves. Children with ASD had a significantly larger percentage of N2 sleep (p=0.003, Cohen’s d=0.87) and lower percentage of N3 sleep (p=0.001, Cohen’s d=0.96) during the first half of the sleep period as well as a lower percentage of REM sleep (p=0.007, Cohen’s d=0.79) during the second half of the sleep period.

**Table 2:**
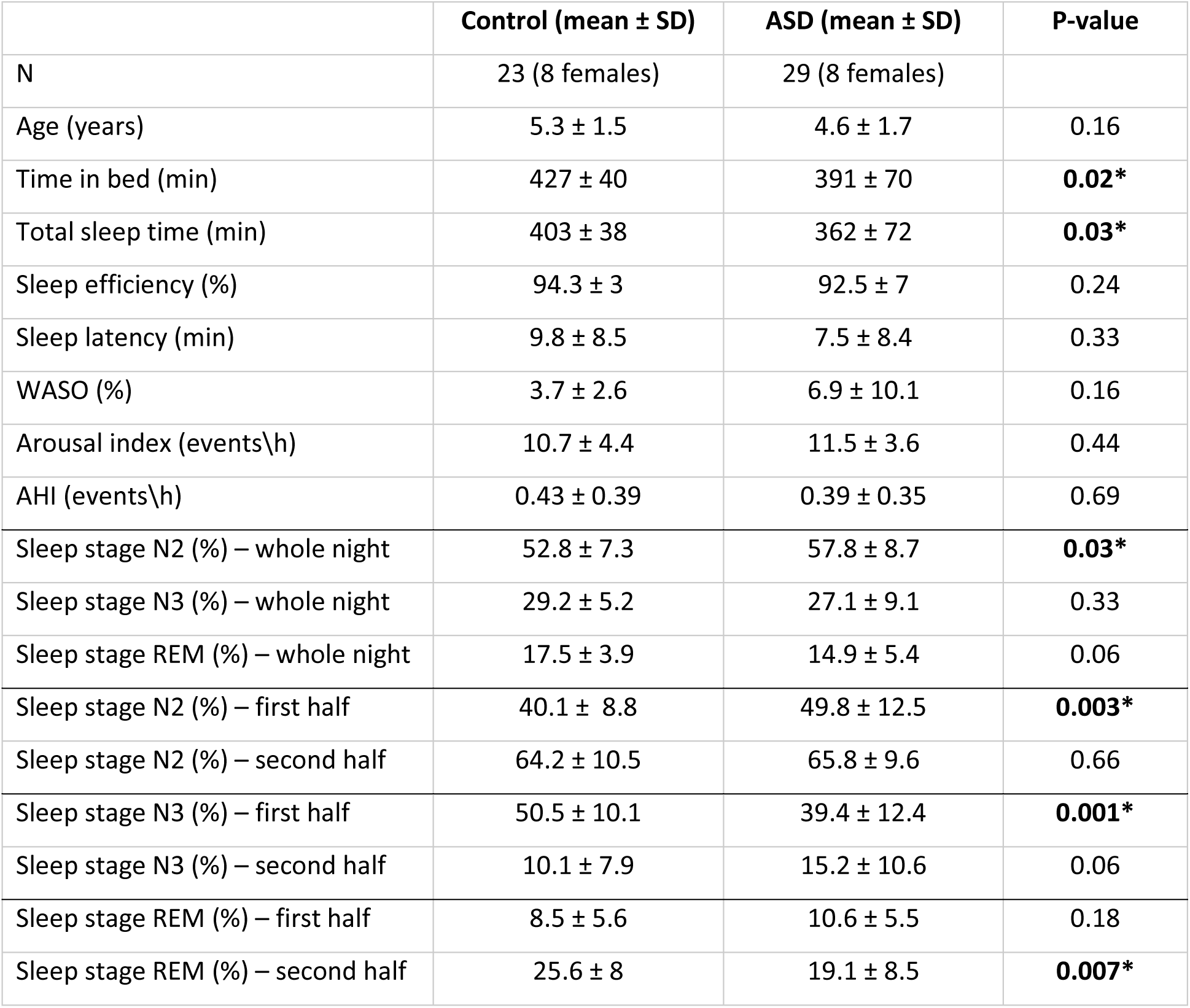
Sleep characteristics in children with ASD and controls. Sleep characteristics in children with ASD and controls. WASO: wake after sleep onset, AHI: apnea-hypopnea index. REM: repaid eye-movement. Asterisks: significant difference as assessed by a two-tailed t-test (p<0.05).

### Differences in Delta power across groups

We computed the spectral power of each frequency band in all artefact-free epochs. We then computed the mean power across all epochs from each sleep stage (i.e. N2, N3 or REM), and compared the findings across children from the two groups. We focused our analyses of EEG power on occipital electrodes given that SWA at the examined ages is maximal in occipital cortex^50^. Children with ASD exhibited significantly weaker power in the Delta (i.e., SWA) and Beta bands during epochs of N3 sleep (Figure 1D). Spectral power in N2 and REM epochs did not differ significantly across groups (Figure 1B&F). Performing the same analyses with the central electrodes did not reveal any significant differences across groups in any of the sleep stages.

**Figure 1:**
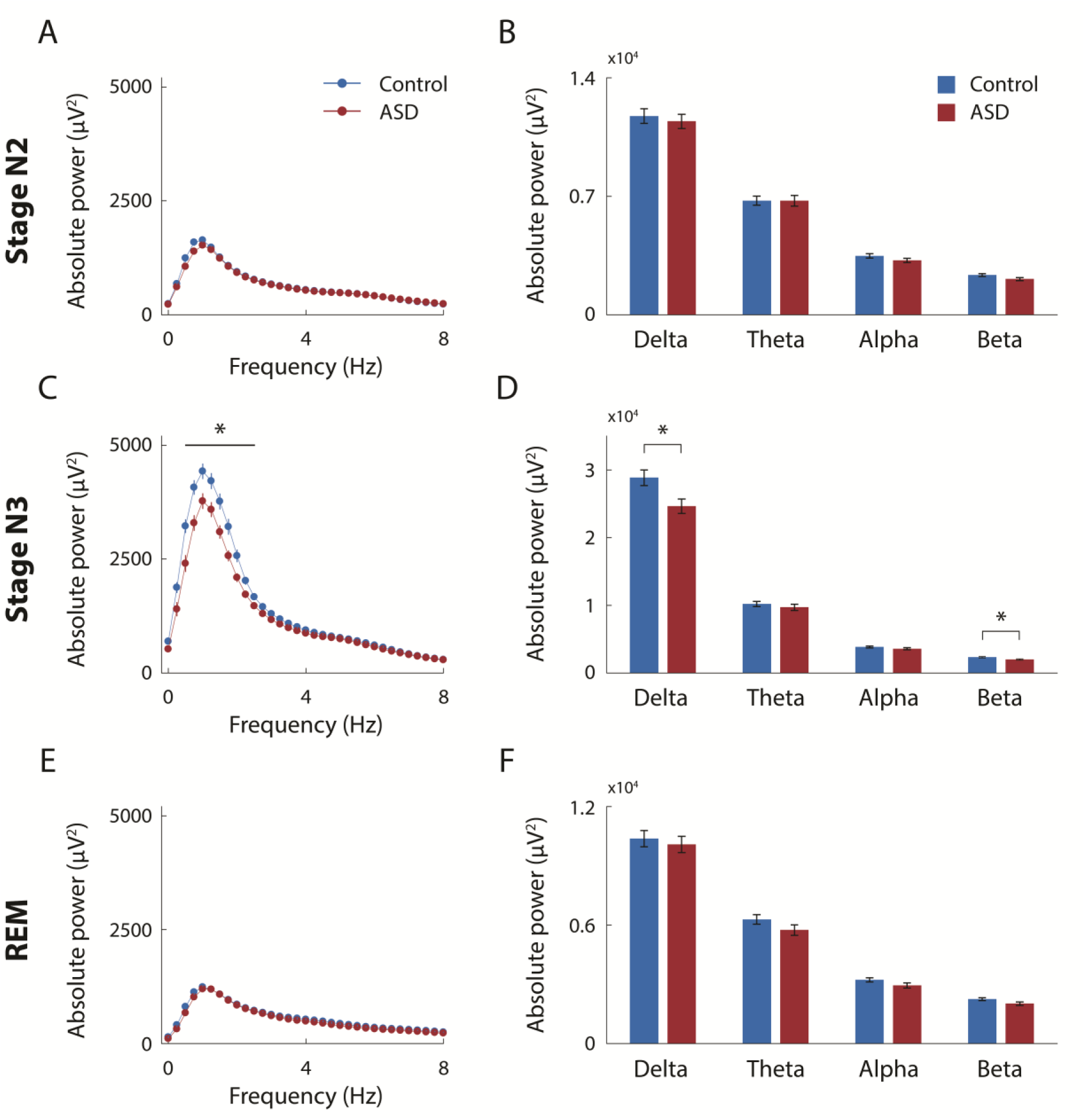
Absolute EEG power for ASD and control groups. Each panel represents the mean power across subjects for the control (blue) and ASD (red) groups during sleep stage N2 (A&B) sleep stage N3 (C&D) and REM sleep (E&F). Power was computed as the mean across all artefact-free epochs from each sleep stage and plotted in 0.25Hz bins (A, C and E) or averaged within the Delta, Theta, Alpha, and Beta frequency bands (B, D and F). Error bar: standard error of the mean across subjects. Asterisks: significant difference across groups (two-tailed t-test, p<0.05, FDR correction).

### SWA dynamics across the night

The power and slope of SWA (i.e., activity in the Delta band, 1-4 Hz) decreased gradually during the night in a manner that corresponded to changes in the proportion of the different sleep stages (Figure 2). These dynamics were similarly apparent in individuals of both groups, yet children with ASD exhibited weaker SWA power particularly in the first two hours of sleep.

**Figure 2:**
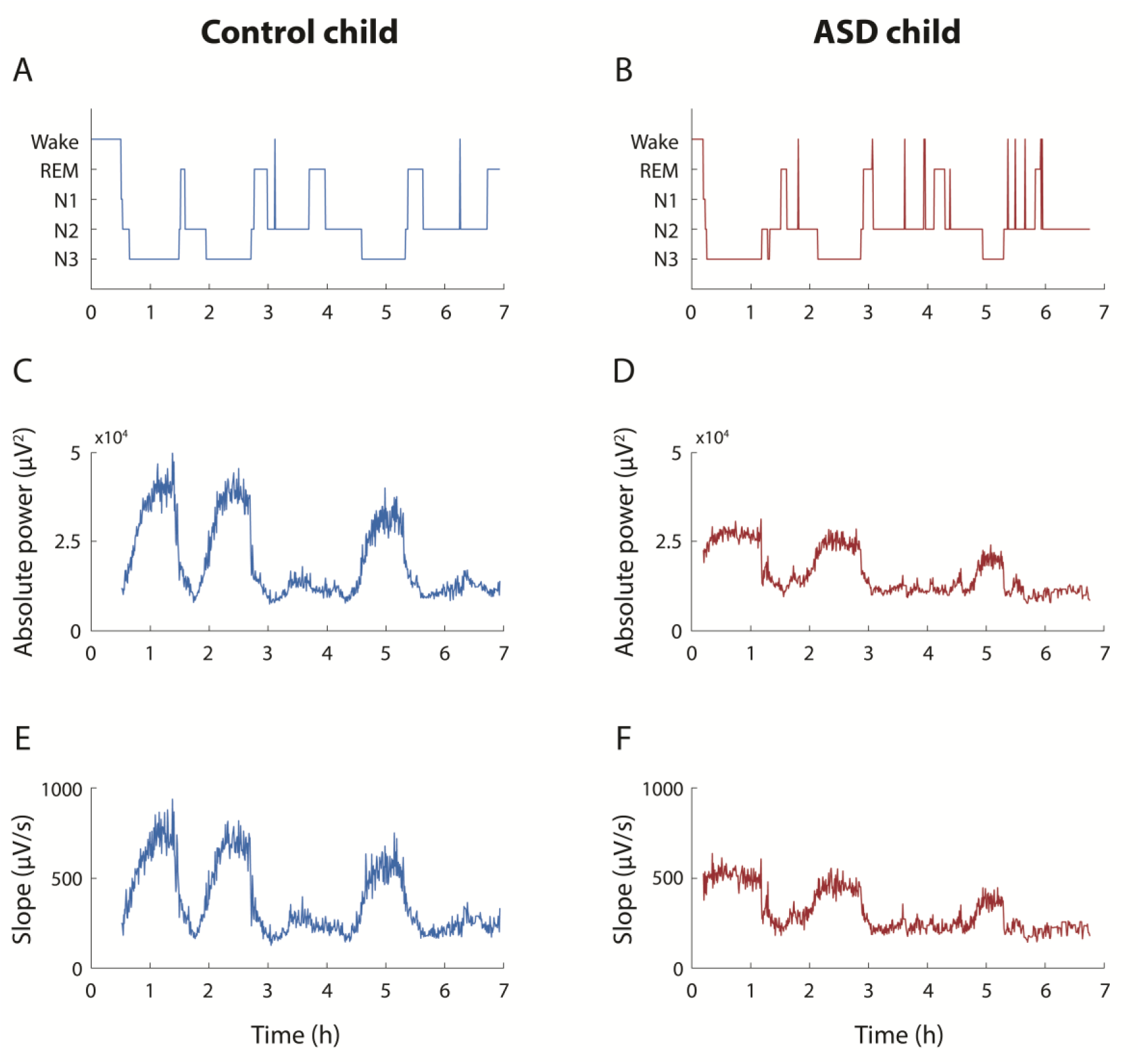
Examples of slow-wave-activity dynamics across the night from one child with ASD (red, right column) and one child from the control group (blue, left column). (A&B) Hypnogram with manual scoring of the different sleep stages in 30 second epochs. (C&D) Time courses of SWA power throughout the night. (E&F) SWA slopes throughout the night. Traces in C-F represent the mean across the two occipital electrodes (O1&O2).

To quantify differences in SWA across groups during specific sleep segments, we divided the night into 1-hour segments, from sleep onset to the end of the PSG evaluation/recording (Figure 3 A&B). SWA power and slope were computed in each 1-hour segment using all non-REM sleep epochs. A two-way ANOVA analysis revealed that SWA power differed significantly across groups (p=0.0008) and over time (p=0.5×10^−30^), with a significant interaction across the two (p=0.02). SWA slope also differed significantly across groups (p=0.001) and over time (p=0.1×10^−29^), with a significant interaction across the two (p=0.02). Follow up comparisons within specific sleep segments using two-tailed t-tests (FDR corrected for multiple comparisons) revealed that children with ASD exhibited significantly weaker SWA power (first hour: p=0.018, Cohen’s d=0.83, second hour: p=0.017, Cohen’s d=0.87) and shallower SWA slopes (first hour: p=0.022, Cohen’s d=0.8, second hour: p=0.022, Cohen’s d=0.85) during the first two hours of sleep. No significant differences were found during the rest of the sleep period. We found similar trends when analyzing data from the central electrodes, but the differences across groups were not statistically significant.

In a complementary analysis we compared the percentage of sleep stages in the same 1-hour sleep segments (Figure 3C) using two-tailed t-tests (FDR corrected). Children with ASD exhibited significantly less N3 sleep during the first hour of sleep (p=0.05, Cohen’s d=0.78), and marginally significant differences during the second hour (p=0.12, Cohen’s d=0.54) as well as a corresponding increase in stage N2 sleep which was marginally significant during the first two hours of sleep (first hour: p=0.1, Cohen’s d=0.63; second hour: p=0.1, Cohen’s d=0.65). Interestingly, ASD children exhibited significantly more REM sleep during the first hour of sleep and a reversed marginally significant trend during the end of the sleep period where they exhibited relatively less REM sleep than controls.

**Figure 3:**
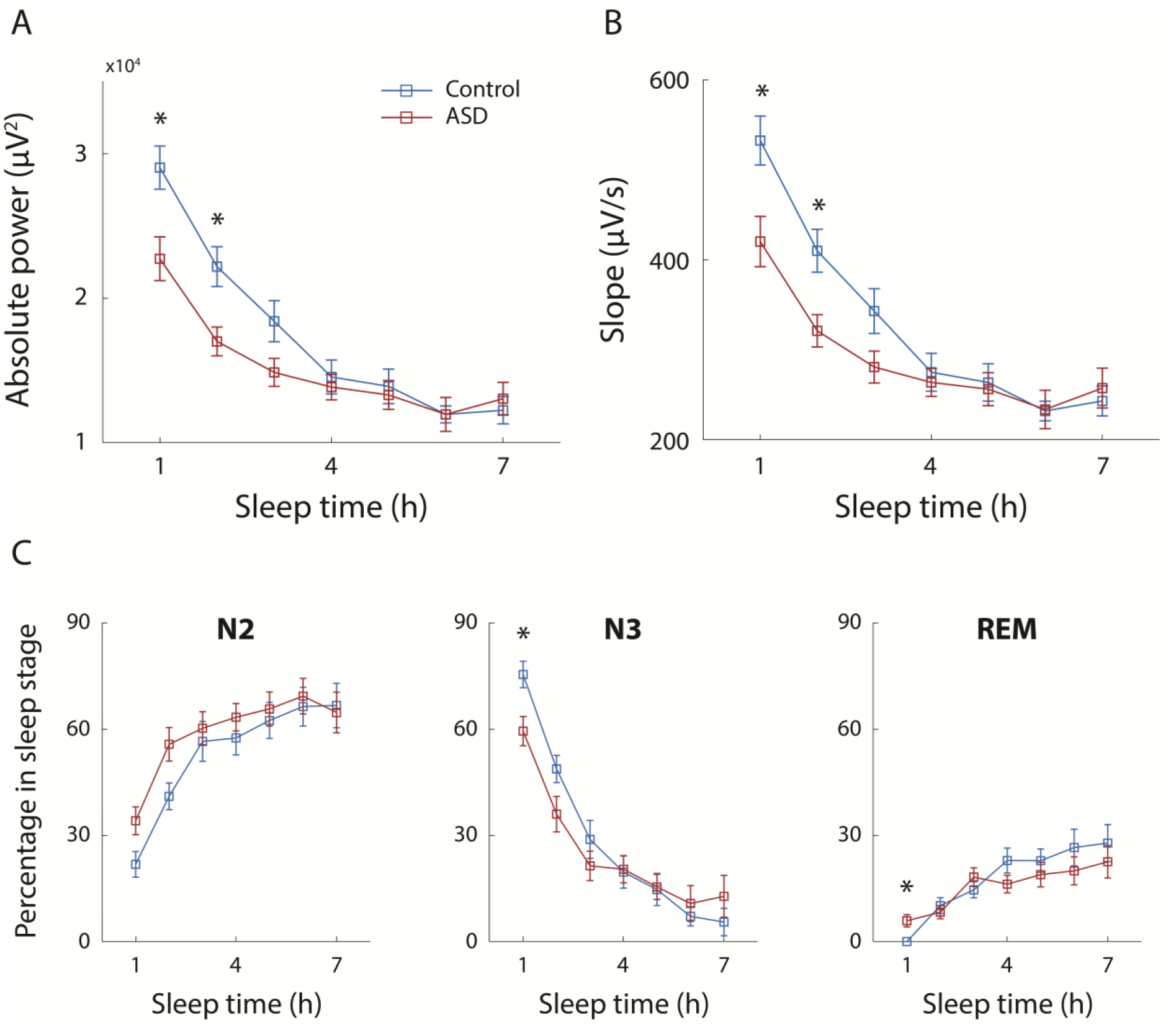
Differences in SWA and percentage of sleep stages across ASD and control groups in 1-hour segments from sleep onset. Power (A) and slope (B) of SWA were computed for each hour of the sleep period, starting at sleep onset, for children with ASD (red) and controls (blue). Percentage of each sleep stage were also computed for each hour (C). Error bars: standard error of the mean across subjects. Asterisks: significant differences across groups (two-tailed t-test; p<0.05, FDR correction).

Previous PSG studies in children and adolescents have reported that SWA power increases with age from early childhood to adolescence and then declines throughout adulthood ^34,51^. To control for differences in the power and slope of SWA, which may arise from differences in the mean age of the control and ASD groups, we performed an identical analysis with a subset of children who were tightly matched for age. This included 21 children with autism (mean age: 5 ± 1.5 years) and 21 controls (mean age: 5 ± 1.3 years). A two-way ANOVA analysis revealed that SWA power differed significantly across groups (p=0.0001) and over time (p=0.2×10^−24^), with a marginally significant interaction across the two (p=0.06). Similarly, SWA slope differed significantly across groups (p=0.0002) and over time (p=0.2×10^−23^), with a significant interaction across the two (p=0.05). Follow up comparisons of each 1-hour sleep segment using two-tailed t-tests (FDR corrected for multiple comparisons) revealed that children with ASD exhibited significantly weaker SWA power (first hour: p=0.034, Cohen’s d=0.78; second hour: p=0.035, Cohen’s d=0.83; third hour: p=0.035, Cohen’s d=0.78) and shallower SWA slopes (first hour: p=0.033, Cohen’s d=0.87; second hour: p=0.033, Cohen’s d=0.84; third hour: p=0.034, Cohen’s d=0.79) during the first three hours of sleep (Figure 4 A&B).

In 15 children with ASD and 4 controls, PSG recording began after the children fell asleep due to difficulties in connecting the EEG electrodes before sleep onset. As a result, a certain period of sleep was missing from the recordings of these children. To control for this potential bias, we performed identical analyses with the 14 ASD children (mean age: 5.6 ± 1.8 years old) and 19 controls (mean age: 5.6 ± 1.4 years old) whose recording began before they fell asleep (Figure 4 C&D). A two-way ANOVA analysis revealed significant differences of both SWA power and slope across groups (power: p=0.003; slope: p=0.004) and over time (power and slope: p=0.8×10^−22^) with a marginally significant interaction across the two factors (power: p=0.064; slope: p=0.065). Subsequent comparisons using two-tailed t-test revealed significantly weaker SWA power and shallower slopes in children with ASD during the second hour of sleep (power: p=0.01, Cohen’s d=1.2; slope: p=0.012, Cohen’s d=1.2) and a marginally significant difference during the first hour of sleep (power: p=0.08, Cohen’s d=0.84; slope: p=0.09, Cohen’s d=0.82).

**Figure 4:**
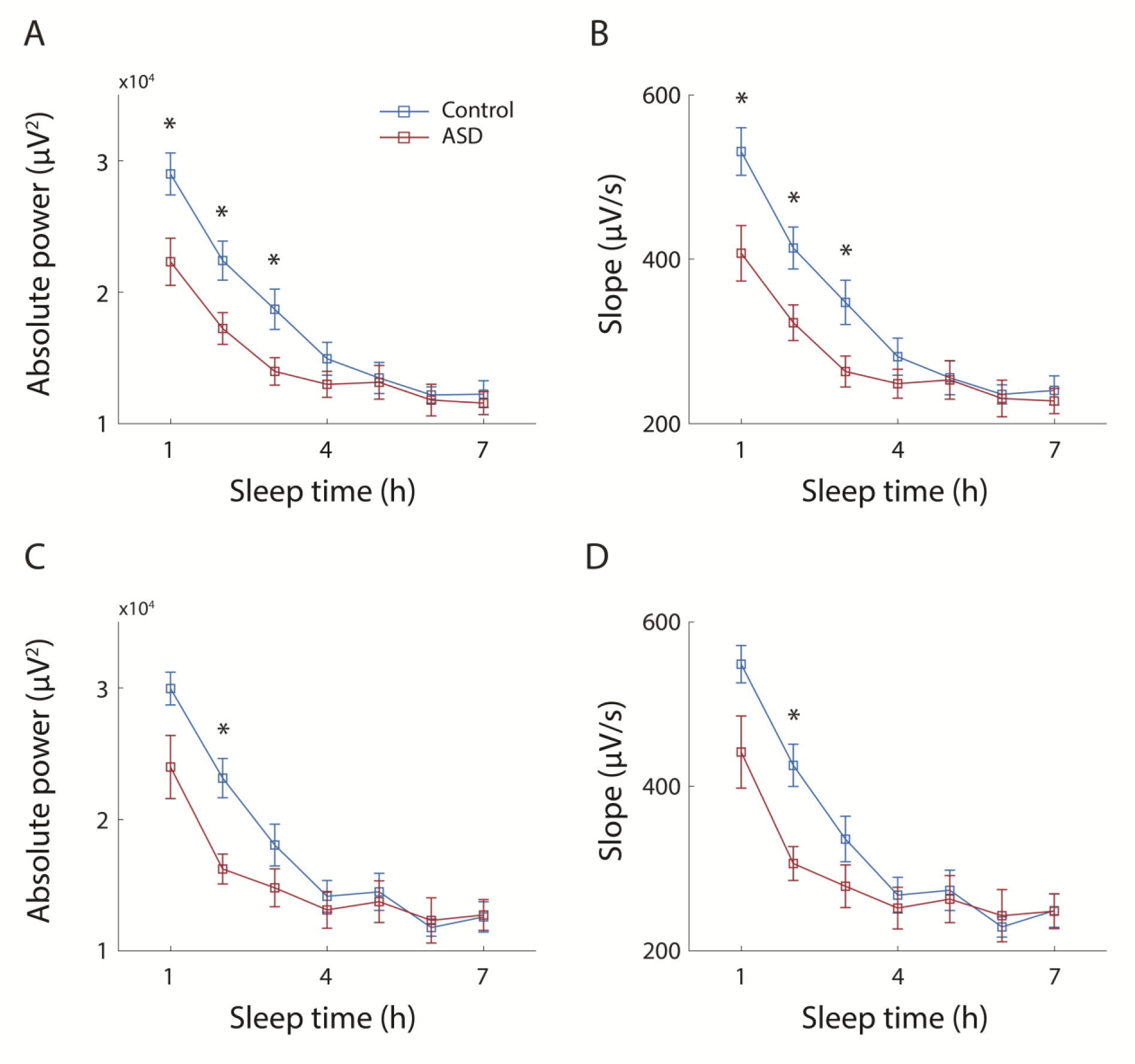
Differences in SWA across groups when tightly matching the children’s age (A&B) and when including only children whose PSG recording started before sleep onset (C&D). Power (A&C) and slope (B&D) of SWA were computed for each hour of the sleep period, starting at sleep onset, for children with ASD (red) and controls (blue). Error bars: standard error of the mean across subjects. Asterisks: significant differences across groups (two-tailed t-test; p<0.05, FDR corrected).

### Relationship between SWA and sleep disturbances in children with ASD

We computed the correlation between SWA power in the first hour of sleep and the behavioral sleep scores that parents reported using the CSHQ. Note that this analysis examines the relationship between a single night of sleep in the lab and the parent reports of sleep problems at home. A significant negative correlation was found between bedtime resistance and SWA power (r(26)=-0.49, p=0.01; Figure 4B) as well as a significant negative correlation between total sleep disturbances and SWA power (r(26)=-0.38, p=0.05; Figure 4A). Furthermore, there was a significant negative correlation between the absolute time that children fell asleep in the lab and SWA power (r(29)=-0.42, p=0.02, Figure 4D) as well as a negative non-significant trend between the child’s sleep onset time at home (according to parent report) and SWA power (r(26)=-0.31, p=0.11, Figure 4C). These results suggest that weaker SWA power in children with ASD at the beginning of the night can partially explain individual problems of initiating sleep, resulting in late sleep onset.

**Figure 5:**
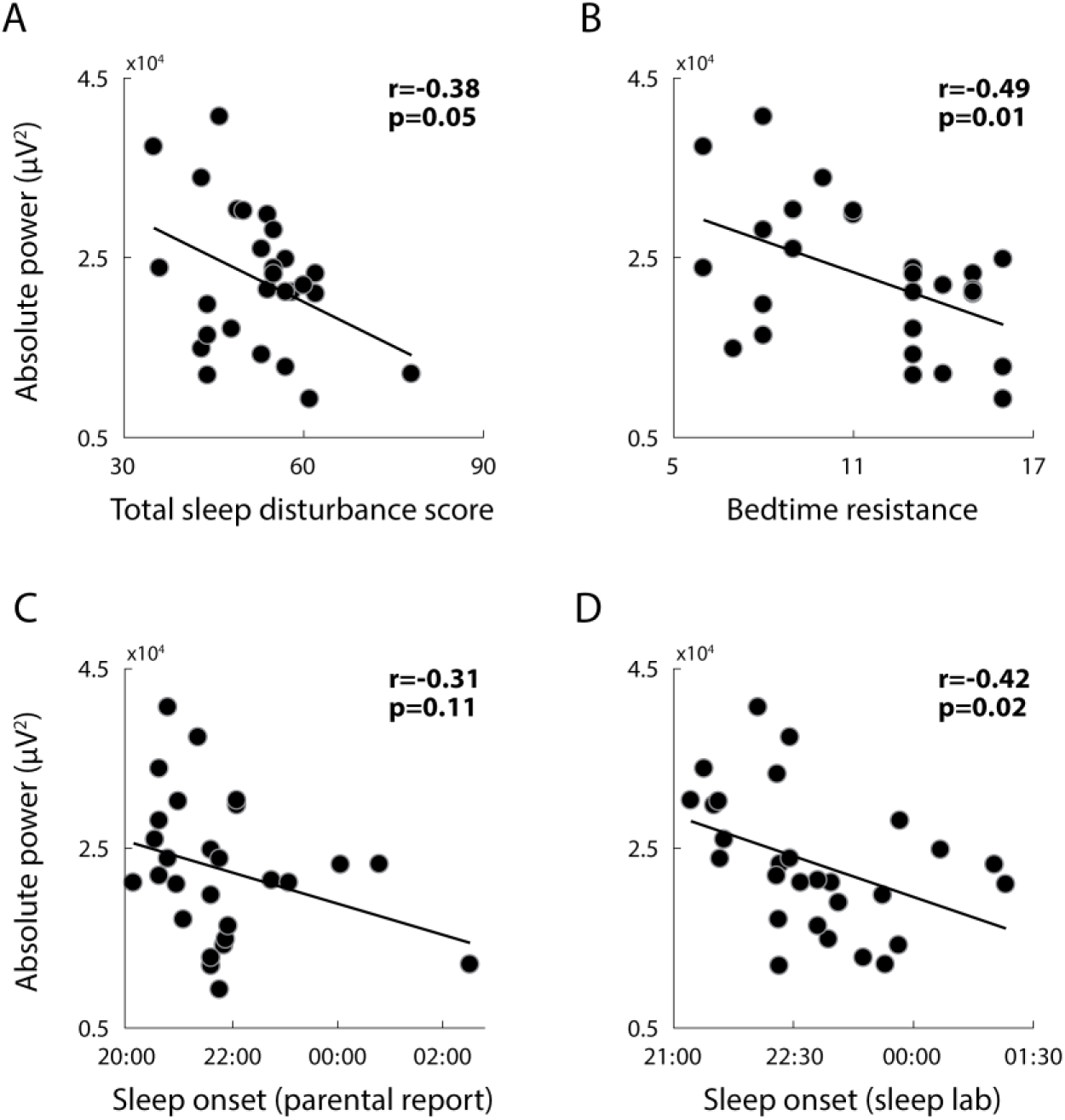
Relationship between SWA power during the first hour of sleep and (A) total sleep disturbance score (B) bedtime resistance score (C) parental report of sleep onset time at home and (D) sleep onset time in the sleep lab. Each point represents a single subject. Pearson’s correlation coefficients and p-values are noted in each panel.

## Discussion

Our results reveal that a considerable number of young children with ASD exhibit weaker SWA power, shallower SWA slopes, and less N3 sleep during the first two hours of sleep (Figures 3&4). We interpret these findings as evidence for a reduction in the pressure to sleep in the ASD children. Moreover, SWA power during the first hour of sleep was significantly correlated with the severity of individual sleep disturbances and especially with sleep-onset difficulties (Figure 5). We, therefore, suggest that a disruption in sleep homeostasis may reduce sleep pressure in children with ASD and exacerbate difficulties with sleep initiation and sleep maintenance.

### Sleep disturbances in children with ASD

Clinical sleep disturbances are apparent in the majority of ASD cases^2,3,5^. While some have reported that sleep problems are more common in children with lower IQ^52^ or higher autism severity^21,52^, others have not^16,20^. More consistent reports have shown that sleep problems are associated with increased self-injury, anxiety, and aggression^18–21^ as well as with sensory sensitivities^15–18^, thereby generating considerable challenges and difficulties for ASD children and their families^53^.

Previous studies have proposed that these sleep disturbances are caused by heightened levels of anxiety^21^, poor sleep hygiene^29^, abnormalities with the melatonin system (i.e., circadian rhythm)^30^, and obstructive sleep apnea (OSA)^31^. Our results suggest that disrupted sleep homeostasis may further contribute to sleep disturbances in children with ASD who do not have OSA. Note that similar disruptions of sleep homeostasis have also been proposed to underlie sleep problems in other populations with insomnia, such as aging adults^54^.

In particular, our results reveal that quantifying SWA power during the first hour of sleep is informative for identifying children with reduced sleep pressure. Like children with OSA who are commonly identified with overnight PSG evaluations, and can benefit from effective targeted treatments^55^, weak SWA during the first two hours of sleep may act as an indicator of children with ASD who may benefit from specific interventions with behavioral and pharmacological treatments that can increase the pressure to sleep^54,56^.

### Quantifying SWA and percentage of sleep stages in children with ASD

We believe that previous PSG studies in children with ASD have not reported the reduced SWA described in the current study for several reasons. First, previous studies did not quantify the amplitude of SWA, but rather relied on manual sleep staging, which is a categorical measure of sleep depth with a very limited range. Our results showed that differences across groups were larger and clearer when quantifying SWA (Figure 3). Nevertheless, one may expect that reduced SWA would be apparent in smaller proportions of SWS (stage N3). This was indeed reported by some PSG studies^22–25^, but not others^26–28^. Note that in our results, the proportion of SWS was not significant different across groups when examining the entire sleep period (Table 2). It was only after we split the night into 1-hour segments following sleep onset, that considerable differences in the proportion of SWS emerged in the first two hours after sleep onset (Figure 3&4).

This emphasizes a second key point: the importance of quantifying SWA and percentage of sleep stages in separate segments relative to sleep onset rather than averaging the measures across the entire sleep period. SWA power and the percentage of sleep stages change dramatically throughout the night^32^ (Figure 2). Meaningful abnormalities in sleep regulation may, therefore, appear in specific sleep segments and be obscured by averaging measures throughout the entire sleep period. Moreover, since children with ASD often sleep less during PSG evaluations^28^, the relative proportion of sleep stages across the entire night will be biased by their shorter sleep duration, which varies across studies.

Taken together, these findings motivate using quantification of SWA power, rather than traditional sleep staging, and focusing on the first hour following sleep onset. This measure is not biased by potential differences in overall sleep duration during the PSG and represents an objective and quantitative marker of initial sleep pressure in individual children.

### The importance of deep slow wave sleep (SWS)

Contemporary sleep research highlights the importance of SWS for regulating the strength (i.e., the number and efficacy) of cortical synapses^36,38,57,58^, which is critical for proper cognitive function including learning and memory consolidation^34,38,59–61^. In typically developing individuals, the amplitude of SWS (i.e. SWA power) increases as a function of time spent awake as demonstrated by studies with sleep deprivation^32^. A remarkable finding in our study is that children with ASD did not exhibit this canonical relationship. Indeed, children with ASD who fell asleep later in the night exhibited weaker SWA (Figure 5). This suggests that the ASD children with the larger sleep onset disturbances had greater sleep homeostasis impairments, and these particular children are likely to benefit most from targeted therapy.

### Limitations

An important limitation of the current study is that the ASD and control samples were not recruited in an identical manner. The ASD children were recruited prospectively from a clinical cohort of a large tertiary medical center and referred by the research team to PSG regardless of potential sleep concerns (see Table 1). In contrast, the control PSG recordings were extracted retrospectively from the same tertiary medical center and included typically developing children who were referred to PSG with sleep concerns. For the current study we only selected PSG recordings of control children who were referred with minor sleep concerns (mostly snoring problems) and completed the PSG without any clinical findings. Nevertheless, it would be important to validate the reported findings with prospectively recruited control children who do not have any sleep concerns.

## Conclusions

The children with ASD who participated in the current study exhibited reduced SWA, which we interpret as an indication of weak sleep pressure. Moreover, the magnitude of this reduction was associated with individual severity of sleep disturbances and particularly with sleep-onset difficulties. A variety of existing behavioral and pharmacological interventions are available for enhancing sleep pressure (i.e., increasing sleep depth)^54^ including mild interventions such as increased exercise^62^. Since improving sleep quality is likely to reduce aberrant behaviors^18–21^ in children with ASD and reduce parental stress^53^, the initiation of clinical trials with these interventions is highly warranted.

## Acknowledgments

This work was supported by a SFARI explorer grant 439370, Israel Science Foundation grant 961/14, and an Israel Academy of Sciences Adams Fellowship to A.A.

